# Evolving learning state reactivation and value encoding neural dynamics in multi-step planning

**DOI:** 10.1101/2025.10.22.683897

**Authors:** Leo Chi U Seak, Raymond J. Dolan

## Abstract

Planning in value-based decision making is often dynamic, with reinforcement learning (RL) providing a powerful framework for investigating how value and action at each step change across trials. Surprisingly, the evolving neural signatures of value estimation and state reactivation in multi-step planning, both within and across trials, have received little consideration. Here, using magnetoencephalography (MEG), we detail neural dynamics associated with planning, wherein subjects were tasked to find an optimal path in order to maximise reward. Behavioural evidence showed improved performance across trials, including subjects showing an increasing disregard for low-value states. MEG data captured evolving value estimation signals such that, across trials, there was an emergence of stronger and earlier within trial value encoding linked to boosted vmPFC activity. Value encoding signals showed a positive correlation and individual performance metrics, as reflected in overall task-related reward earnings. Strikingly, across trials, there was an attenuation of state reactivation for negative-value states, an effect that positively correlated with evolving negative-value state avoidance behaviour. The finding linking neural dynamics, including a valence-dependent selective reactivation of negative states, to across-trial behavioural improvement advances an understanding of learning during multi-step planning.

## Introduction

Acquired value is a dominant guiding force in decision making (Bhatia et al., 2018). When buying groceries, navigating to new places, and engaging in long-term planning, we rely on estimates of learnt values (Rangel et al., 2008). Our decision making is also dynamically configured, both within and across decision steps, such that consequences at one step significantly impact the choices we make at the next (Dayan, 1993; Cisek et al., 2009; Ullsperger et al., 2014; Keramati et al., 2016; Miller & Venditto, 2021; Zylberberg et al., 2024).

In multi-step planning, tree search (Daw et al., 2005), successor representation (Momennejad et al., 2017), dynamic programming (Mattar & Daw, 2018), and hierarchical reinforcement learning (RL) (Eckstein & Collins, 2020) have variously been invoked to explain state value estimation and actions. Common to these is a reinforcement learning (RL) framework that addresses how agents interact with an environment, invoking value estimation, action planning and outcome-dependent value updating (Sutton & Barto, 1998; Dolan & Dayan, 2013; Miller & Venditto, 2021). In simple terms, agents learn the value of states and associated actions across multiple trial and error iterations.

To reduce the computational burden of multi-step planning, humans and RL agents learn to disregard action states with low or negative value (Huys et al., 2012; Momennejad et al., 2017; Faulkner et al., 2021). Therefore, a major challenge is a characterisation of the underlying state representation dynamics associated with this type of value learning (Hunt et al., 2021). Recent rodent studies suggest a value-dependent state reactivation of place cells (Jin et al., 2024), while in humans, state reactivation of chosen paths is reported to be stronger than for unchosen paths (Wise et al., 2021). Nevertheless, how the state reactivation signals evolve in accordance with value learning in multi-step planning remains under investigated.

Planning-related neural activity can be considered from the perspective of within and across-trial dynamics, respectively. Prior studies of within-trial value dynamics have variously addressed credit assignment (Hamid et al., 2021), arithmetic computations (Pinheiro-Chagas et al., 2024) and reward attributes (Hunt & Hayden, 2017; Hunt et al., 2018; Grabenhorst & Báez-Mendoza, 2025) with dynamics elucidated using approaches that include drift-diffusion models (DDM) and attractor network models (Ratcliff 1978; Hopfield 1984; Ratcliff et al., 2016; Urai et al., 2019; Prat-Ortega et al., 2021; Zylberberg et al., 2024; Ging-Jehli et al., 2025). By contrast, studies of across-trial dynamics have focused on associative learning (Biane et al., 2023; Kim et al., 2024) and adaptation to dynamic sensory inputs (Ruesseler et al., 2023). Notably, a number of these studies have also addressed how value learning signatures evolve within and across trials (Gluth et al., 2012; Hunt et al., 2015; Ebitz et al., 2018; Collins & Frank, 2018; Yoo & Hayden, 2020; Miller et al., 2022). Despite increasing recognition of its importance, the question of how state (and value) dynamics evolve during multi-step planning and learning remains largely unexplored.

Here we used Magnetoencephalography (MEG) to investigate the dynamics of value learning during multi-step planning, including how state reactivation and value-encoding signals evolve over the course of learning. Subjects were tasked to perform a 4-step, non-spatial, navigation task in order, in each trial, to maximise monetary reward. During planning, we found an emergence of stronger and earlier across-trial value encoding signals, linked to increased vmPFC activity. Across-trial learning was also linked to an increasing preferential reactivation of non-negative states.

## Results

### A task to measure planning

To address how humans learn multi-step planning, we availed of data from a study design (Kurth-Nelson et al., 2016) wherein participants performed a 4-step, non-spatial, planning task (Fig. 1A & 1B). Participants completed this task while undergoing MEG brain scanning. Our prior work showed that MEG can detect content-rich neural state reactivations, both during planning and at rest, even when the state in question is represented without concurrent sensory input (Liu et al., 2019; Nour et al., 2021).

**Figure 1.**
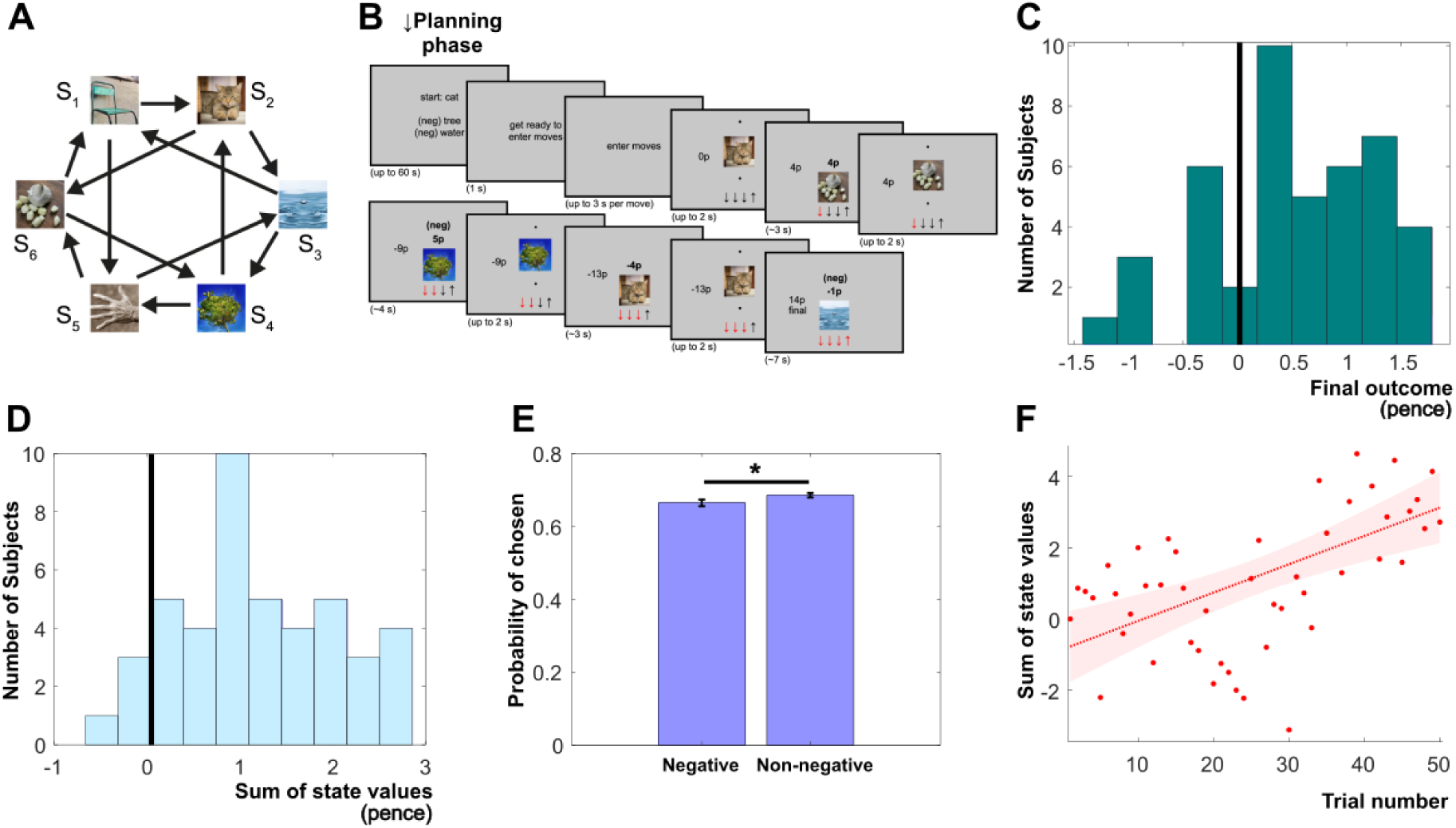
Experimental design and behavioural data. Participants were tasked to maximize a final reward through multi-step, non-spatial, navigation. At each trial, a starting state was pseudorandomly assigned, and participants were tasked to enter four moves. (A) The diagram illustrates the transition structure between six available states, each defined by a distinct visual stimulus. From any occupied state, the participant could choose to navigate to one of two linked states. A drifting value was assigned to each state such that, to maximise reward, participants needed to vary their planned moves across the course of the experiment. (B) At the start of each trial, participants were informed of their starting state as well as the trial assigned two “neg-flip” states. Navigating to the latter would result in the accumulated reward flipping sign (from positive to negative or vice versa). Participants were allowed to plan for up to 60 seconds, and then then allowed 3 seconds to enter their chosen moves without feedback. Next, the chosen sequence was played out such that subjects would see their chosen visual stimuli (states), the reward value of each state and their accumulated reward. (C) Histogram distribution of participants’ performance averaged within each participant (significant above chance, signed-rank test against 0: p=7.6067e-05). X-axis indicates the performance (pence they would attain if the states had not changed since the last time they saw it). Y-axis indicates the number of participants. (D) Histogram distribution of participants’ summed chosen state values in pence (for each trial, the sum of the 4 states (steps); averaged across trials). Significant above chance performance was found (signed-rank test p=3.6187e-08 against 0). (E) Bar chart showing averaged probabilities of choosing negative states versus non-negative states (baseline: 4 chosen states out of the 6 total states or 66.66%), in which more non-negative states were chosen. Error bars showing mean ± SEM across sessions (N=44). * denotes p<0.05, two-tailed Mann–Whitney U test. (F) Participants’ disposition to disregard low-value states, evident in an increasing choice of higher value states (summed chosen state values) across the trials (significant Spearman correlation with rho=0.5854, p=7.99e-06, N=44). Dot represents the averaged value (Y-axis) across participants for the trial (X-axis). The red line represents the least-squares line of all the dots and the shaded region represents the 95% confidence interval. A and B were adapted from Kurth-Nelson et al., 2016.

Within the task, at each trial, participants were randomly assigned one of six possible starting states (S1-S6 in Fig. 1A) and then entered four sequential moves. At each state, this entailed one of two manually entered move options (“up” and “down”) that led deterministically to a successive state. For example, at state S1, a participant could choose to go to either S2 or S5. Each state was defined by a unique visual image and linked to a pre-determined reward amount (that varied across states from -5 to 5 pence), with the latter drifting pseudo-randomly across trials by amounts of 1p, 0p, or 1p respectively.

Prior to MEG scanning, to ensure participants understood the task structure, they were familiarised with the general structure of the task, but not the value associated with each state. They were also instructed to plan ahead over a horizon of four steps with an explicit goal of harvesting as much reward as possible. Prior to progressing to the scanning stage, they were required to attain 100% accuracy on a quiz that probed their knowledge of the task transition structure.

Our key question pertained to neural state reactivation dynamics during planning, both within and across trials, as detected using MEG. Participants were allowed up to 60s (self-paced) to plan a sequence of moves before entering them on a keyboard. This variable planning phase, prior to move entry, provided the primary context for our MEG analyses of neural dynamics. To encourage subjects to plan each of the steps, and to minimise the possibility of relying on a simple stimulus-response learning strategy, two (pseudorandomly selected) “neg-flip” states were introduced at each trial, such that when participants reached that state, their cumulative reward was multiplied by -1.

### Behavioural data

We first assessed task performance, based on what subjects earned (value they would get if states had not changed since last encountered, equivalent to having a 1.0 learning rate; see Methods for details). Overall, we found that across 44 sessions, there was above-chance performance in 33 of 44 sessions (averaged final outcome). Average performance (Fig. 1C) across all 44 sessions was significantly above chance (0), based on a signed-rank test (p= 7.6067e-05). Consistent with our previous findings (Kurth-Nelson et al., 2016), we found a weak, albeit significant, correlation (significant Pearson Rho = 0.0440, p=0.0383, but not Spearman Rho = 0.0345, p=0.1044) between reaction time and enhanced performance.

Next, we investigated if participants were disposed to choose fewer low-value states, as observed in previous decision planning studies (Huys et al., 2012; Faulkner et al., 2021). To test this, we summed values (values based on last-seen, as detailed above) for the 4 chosen states (steps), within each trial, for each participant. Across participants, the sum of chosen state values (averaged across trials) was significantly above zero (Fig. 1D; signed-rank test p=3.6187e-08), indicating participants were more disposed to choose higher-value states than lower-value states. We also calculated the probability of choice of negative states and non-negative (including positive and zero) states, for each trial and each participant. Averaging across trials, we found that overall participants chose more non-negative states than negative states (Fig. 1E; N=44, Mann–Whitney U test p=0.0389).

To test if participants learnt to selectively disregard low-value states across trials, we ascertained if, over the course of trials, participants showed an increased propensity to choose states with higher values. We first calculated, for each trial and each participant, the sum of the 4 chosen state (step) values (values based on last-seen, as mentioned above) and correlated these with trial number. We found positive correlations in 40/44 sessions, with all 18 out of 44 significant correlations being positive (Fig. 1F & Fig. S1). There was also a moderately strong Spearman correlation (Rho=0.5854, p=7.9869e-06) when averaging across all participants, indicating participants chose fewer low-value states across trials.

### Neural dynamics of value estimation: learning of final outcome value

Next, we asked if neural dynamics reflect learning of value during planning. To index this, we used Generalised Linear Models (GLM) to predict a final trial reward amount (i.e. the value they would get if states had not changed since last encountered). Because of diverse planning times, across trials and across participants, we down-sampled the sensor-based MEG data to 10 time points for each planning period and then input these into a GLM. To ask whether an evolving neural signature reflects value learning, we divided trials into 4 quartiles based on trial number (i.e. the period of time in the experiment), an approach akin to that implemented in recent animal studies (Amo et al., 2022; Ren et al., 2025). We then built a separate GLM for each quartile per sensor, averaged across sensors, and represented value encoding signals (GLM coefficients) as a heatmap (Fig. 2A). In effect, these GLM coefficients represent value encoding signals at each time point in a trial.

**Figure 2.**
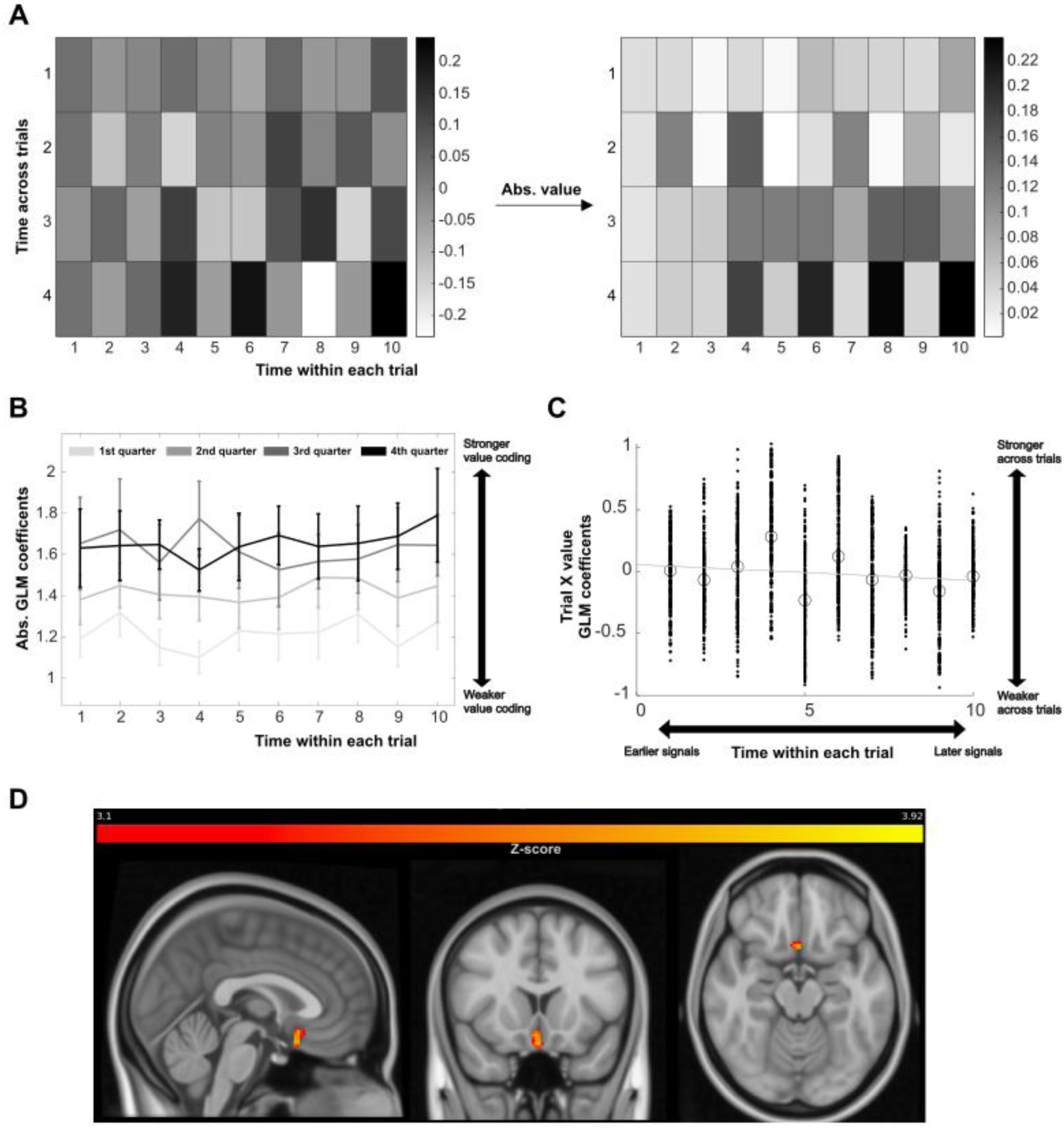
GLM analyses of value-encoding neural dynamics. Dynamics of value neural encoding during planning. To account for diverse planning times, MEG data were first down-sampled to 10 time points. We used a GLM to predict the final outcome value a participant obtained for each trial. To test across-trial dynamics, we built a GLM for each of the 4 quartiles of trials in A-C (e.g. first 25% of trials enter the 1^st^ GLM; the next 25% of trials enter the 2^nd^ GLM, etc). (A) A boost in value-linked activity following learning, as illustrated by heatmaps of GLM coefficients across the 10 time points (X-axis) and across the 4 quartiles (Y-axis). The heatmap on the left shows averaged (across sensor and across participant) GLM coefficients within, and across, quartiles. To illustrate overall value encoding, absolute GLM coefficients (right panel; absolute after averaging across sensors) are also plotted. Grey scales represent the GLM coefficients. 2-way ANOVA (both across 10 planning time points and 4 quartiles of trials; p<0.0001) revealed significant neural dynamics of the absolute coefficients within and across trials. Post-hoc Spearman correlation (rho = 0.1377, p = 6.5878e-09) shows a significant increase in value-encoding activity following learning. (B) Value-encoding signal increases across trials, plotted as absolute GLM coefficients signals (absolute before averaging across sensors; this allowed us to get all value-encoding signals without considering if they coded negatively or positively). This shows a significant increase in value-encoding coefficients across each quartile of the experiment (Spearman rho = 0.2159, p<0.0001). Error bars showing mean ± SEM across 44 sessions. (C) Within-trial dynamics based on mixed-effect models. Instead of dividing trials into 4 quartiles, we include trial numbers and the interaction of final reward amount (value they would get if states had not changed since last encountered) and trial numbers as regressors. These coefficients were entered into a group (second) level GLM. Each solid dot in this scatter plot represents a sensor’s GLM coefficients (interaction of value and trial numbers; Y-axis) along the 10 time points (X-axis). The negative correlation (Spearman rho = -0.0942, p = 8.5319e-07) of these two axes showed the emergence of a stronger value-encoding signal towards the beginning of a planning period. Each hollow circle represents the mean GLM coefficients of all sensors at that time point. The grey line represents the least-squares line. (D) Source reconstruction for an a priori ROI showing significantly greater vmPFC activity with learning (Z>3.1, last > first MEG session, permutation n=5000, FWE corrected p<0.05 cluster level significant), using an independent vmPFC mask derived from previous studies (Delgado et al., 2016; Bhanji et al., 2019).

In a 2-way ANOVA, we first asked whether there were changes in value encoding signals reflective of either within (quartile of 10 time points) or across-trial (4 quartiles) dynamics. We found significant GLM main effects for value coding coefficients, both within (10 time points) and across-trial (4 quartiles) (p<0.0001 for both), indicating evolving value encoding neural dynamics.

Next, using an analogous methodological approach to previous human and animal studies (Amo et al., 2022; Huang et al., 2024), we examined the evolution of overall value encoding signals irrespective of whether these were positive or negative, taking the absolute values of the GLM coefficients for each time point and quartile (Fig. 2A, right) and asking how these changed across quartiles. The emergence of neural value encoding signals indicative of learning across quartiles was evident in a significant positive Spearman correlation (rho = 0.1377, p = 6.5878e-09). As these value encoding signals were GLM coefficients (beta estimates), in contrast to evoked potentials, an increase is suggestive of a boost in value-linked activity, as opposed to a mere reflection of non-specific time-related effects.

To confirm learning dynamics across quartiles, we also examined the absolute values of GLM coefficients, without averaging across sensors. In comparison to the above analyses, this approach negates averaging out positive and negative signals between sensors, arguably providing a less biased index of neural activity across quartiles. This revealed a significant increase across quartiles (Spearman rho =0.2159, p<0.0001 ; Fig. 2B), consistent with boosted value encoding signals as a function of learning across quartiles.

We next asked how within-trial dynamics evolve across the course of the experiment, using a mixed effects model with trials, values and their interactions as regressors. We found a significant negative Spearman Rho correlation between time points and trial-value interaction (Fig. 2C; Spearman rho = -0.0942, p = 8.5319e-07), indicating that, over the course of trials, there was an evolution of richer value-encoding signals towards the beginning of a planning period.

Prior investigations of learning-related brain dynamics consistently implicate an involvement of vmPFC (Gläscher et al., 2009; Moneta et al., 2024), leading us to focus on value encoding dynamics within this region. To this end, we implemented a beamforming analysis (1-45Hz), focusing on the first two seconds of the planning phase across trials, using a previously used protocol (Van Veen et al., 1997; Liu et al., 2019; Nour et al., 2021). Comparing the first and last blocks of the experiment, we found increased activity in vmPFC (Fig. 2D; p=0.025; small volume corrected with an out-of-sample mask), consistent with prior human neuroimaging studies (Gläscher et al., 2009; Chib et al., 2009; FitzGerald et al., 2010, 2012; Smith et al., 2010; Rushworth et al., 2011; Boorman et al., 2013), as well as findings from non-human primates (Roberts & Clarke, 2019; Bongioanni et al., 2021; Veselic et al., 2025), that implicate vmPFC in value-based decisions.

The above results highlight evolving temporal neural dynamics reflective of a final reward outcome (value estimation), within and across trials, that in turn are linked to an across-trial increase in vmPFC activity.

### Neural dynamics and selective reactivation of negative states

A feature of value-based decision making is the emergence of a disposition to avoid pathways with negative states (Huys et al., 2012; Faulkner et al., 2021). One hypothesis is that this phenomenon might reflect the strength of neural representation (reactivations) of avoided negative states, akin to selective reactivation reported in rodents (Jin et al., 2024). To identify neural activity patterns linked to each task state, and index these during planning, we employed a neuronal decoding approach validated in prior studies (Kurth-Nelson et al., 2016; Liu et al, 2019). Specifically, for each participant, we trained Lasso-regularised logistic regression models to predict state identity from sensor-level MEG signals. This approach uses MEG data from a functional localiser task phase, wherein participants were presented in turn each task state (stimuli/image) and reported whether the stimulus was upside down (in 20% of trials and they were not used for training). As in previous studies, regression models were trained at 200ms after stimulus onset (Kurth-Nelson et al., 2016; Liu et al, 2019), with these object images shown multiple times in random order. As shown in Fig. 3, these decoders had peak accuracy 20.5% above baseline (Fig. 3A & 3B), including an above chance predicted probability for all six decoded states (Fig. 3C).

**Figure 3.**
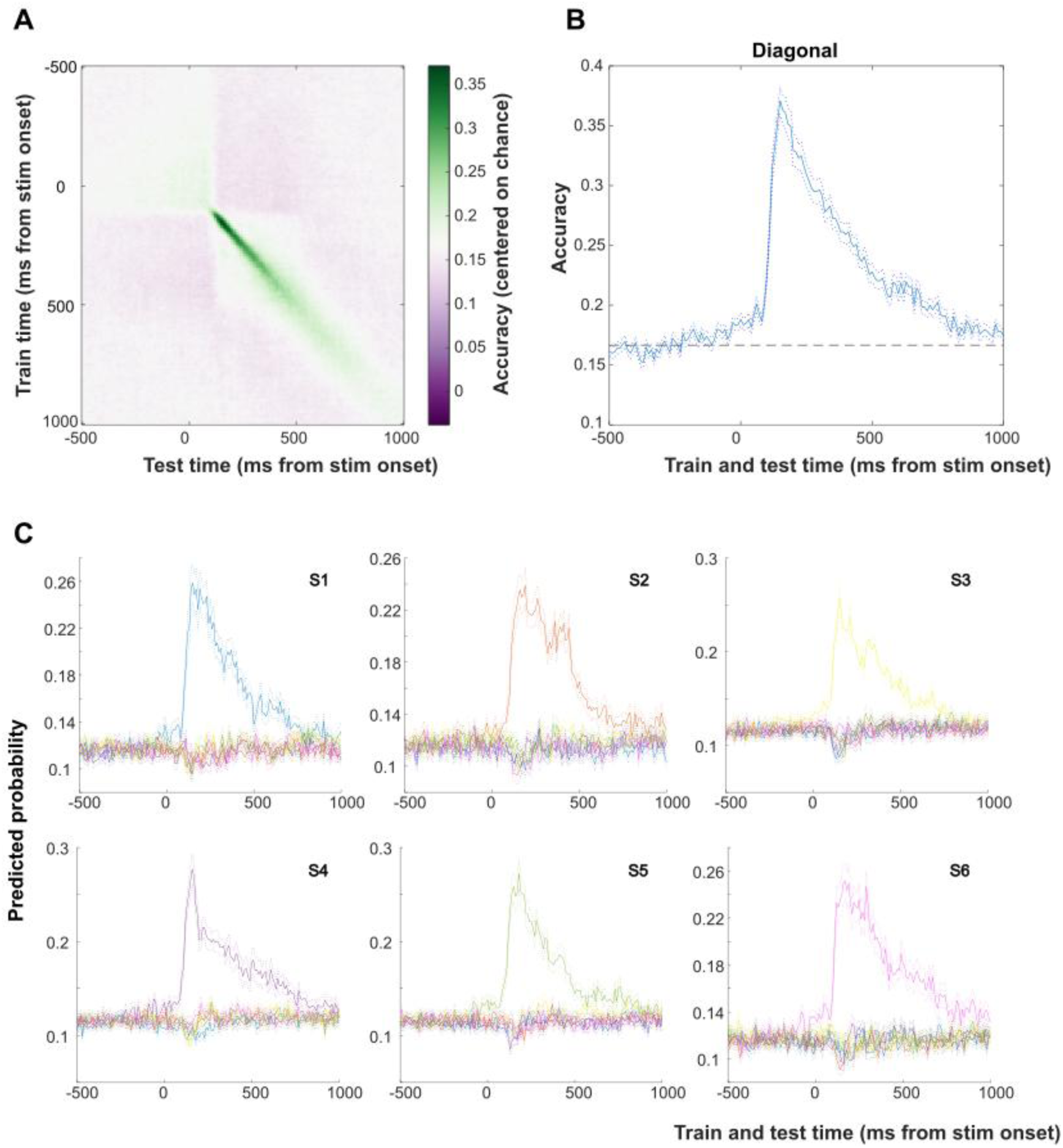
Lasso-GLM decoder performance. For each participant, we built Lasso-GLM decoders to predict states (one of the six) during stimulus onset, when the visual image was shown. To train these decoders, at the end of the experiment (localiser task), we presented these six visual images multiple times and in a random order (see Methods for details). (A) Lasso-GLM decoding accuracy with different training and testing time points. 0 represents the time when stimuli were shown. Green represents above-chance and purple represents below-chance decoding accuracy. A peak decoding accuracy was found at 150ms after stimulus onset (trained and tested at 150ms). (B) Decoding accuracy when trained and tested at the same time point (diagonal line of A). Solid line represents mean, and dotted line represents SEM across 44 sessions. Horizontal dashed line represents decoding at chance level (16.67% or 1/6 states). As in Figure 3A, above chance level decoding was seen after the stimulus onset, with the peak decoding accuracy at 150ms after onset, where the accuracy is 37.2% (20.5% above the baseline). (C) Predicted probability of each of the six states. The ground truth states of the decoders are S1: blue; S2: orange; S3: yellow; S4: purple; S5: green; S6: pink. Each colour line represents one of the six predicted states. High predicted probability peaks were found for all states in accord with ground truth, validating the utility of the decoders. Dotted lines represent the SEM of the solid line (mean) across the 44 sessions.

To ascertain whether neural representations of negative states were disregarded, as predicted based on prior animal data (Jin et al., 2024), we asked whether the decodability strength of negative states during planning was less than that for non-negative (positive & zero) states. Applying trained models to planning phase MEG data, when no actual visual object was shown, for each 10ms bin we ascertained a predicted state reactivation (one of the six possible states). In effect, within a trial, we calculated the decodability of each state and averaged separately across trial decodability of all negative-value and non-negative-value states (baseline decodability: 1/6 or 16.67%). We found decodability of states with negative values was significantly less strong than was the case for non-negative values (Fig. 4A; Mann–Whitney U test: p=0.0494, N=44), mirroring a behavioural avoidance of negative value states (Fig. 1E).

**Figure 4.**
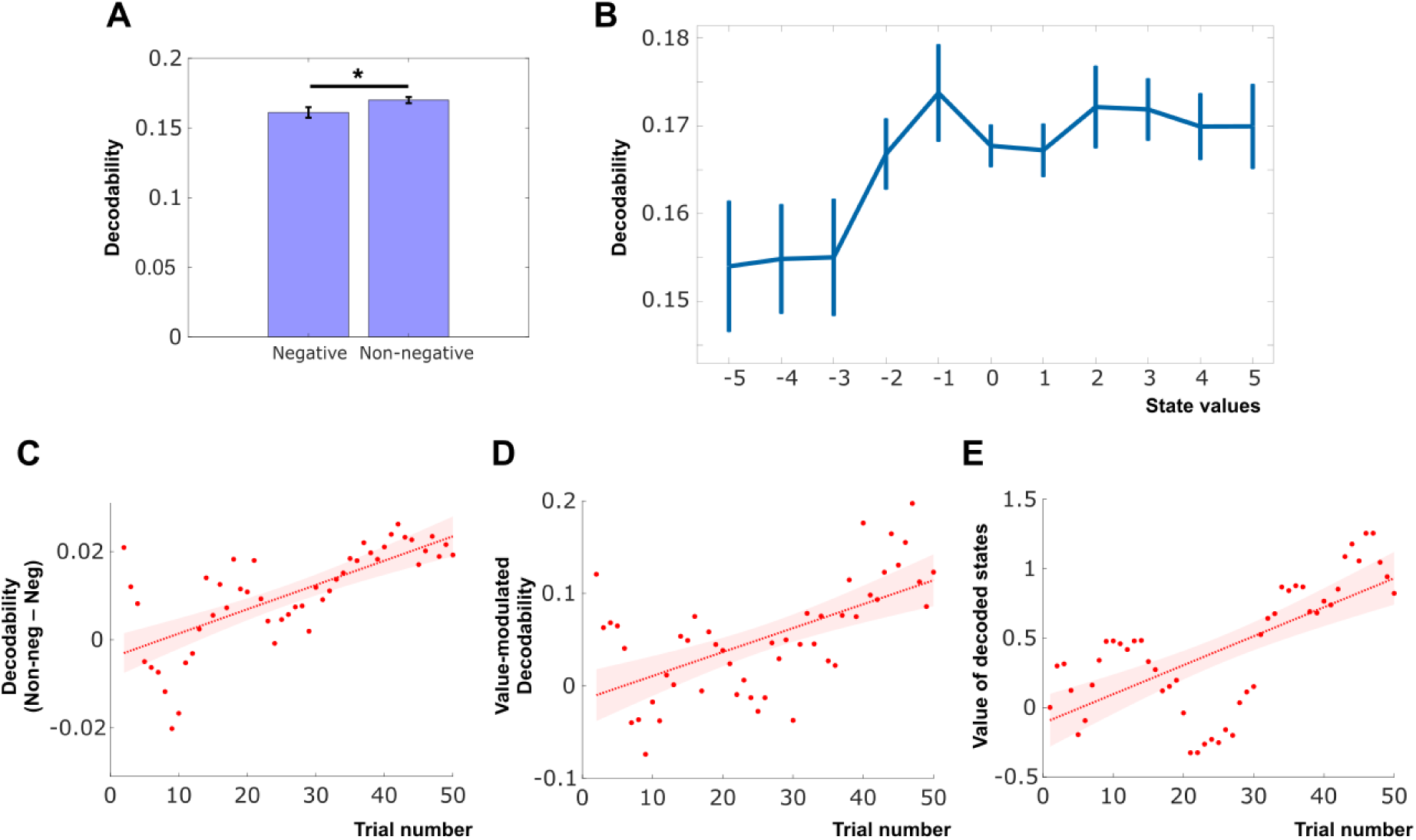
Changes in state decoding following across-trial learning. For each subject, we used trained Lasso decoders of stimuli to predict stimulus identity (one of six possible stimulus states; see Figure 3 for the performance of the decoders). We then applied these trained decoders to MEG activity recorded during the planning phase (decodability; for each of the 10ms). Decodability was averaged across states, including negative/non-negative values (For A), or with all the states that got the same state value (For B). For A &B, we averaged across all the trials; while for C&D, for each trial, we averaged the decodability across participants and plotted these against trial numbers (X-axis). For E, we plotted the sum of the value of the decoded states (Y-axis) against trial numbers (X-axis). (A) Bar chart showing decodability for negative and non-negative value states (the state with maximum decoded probability) during planning. * denotes p<0.05, two-tailed Mann–Whitney U test. Baseline chance level: 16.67% or 1/6, representing a random state was decoded out of the 6 possible states. (B) Line chart (mean ± sem) showing decodability of states as a function of their associated value (the state value shown to the participant last time). Error bars showing mean ± SEM across sessions (N=44). Across negative-value states (−5 to -1, increment by 1), we found a significant positive correlation between state value (X-axis) and decodability (Y-axis), with Spearman Rho=0.0860 and p= 9.9675e-10. When considering all 44 sessions’ correlation Rho: p=0.0266 with significant signed-rank test above 0. Such an effect was not observed for non-negative states (Spearman Rho=0.0134; p=0.2225; signed-rank test for all sessions’ Rho: p=0.8886). These results indicated a positive correlation between state decodability and state value for negative states. (C) Scatter plot showing increase of relative decodability (decodability of non-negative minus decodability of negative states) across trials (Spearman rho = 0.7232, p=2.69e-08, N=44). Plotting separately across-trial (Fig. S2), the effect can be seen to be driven by both an increase in decodability for non-negative states (Spearman rho = 0.7508, p<0.0001) and a decrease in decodability for negative states (Spearman rho = -0.6636, p<0.0001). In C-E, the red lines represent the least-squares lines of all the dots and the shaded regions represent the 95% confidence intervals. (D) Scatter plot showing an across-trial increase in value-modulated decodability (i.e. the trial-by-trial Spearman Rho of state decodability and their associated state value; Spearman rho = 0.5923, p=1.13e-05, N=44). Y-axis represents the averaged Spearman Rho for each trial, across 44 sessions; and X-axis represents the trial numbers. (E) The value of the state being decoded increased across trials (Spearman rho = 0.6640, p=3.70e-07, N=44), consistent with the decodability of low-value states being disregarded following learning in the behaviour (Fig. 1F). For each trial, we summed values of the decoded states, and averaged them across 44 sessions as Y-axis. Such a trial-by-trial value was then tested for correlation with the trial number (X-axis).

Next, we tested if selectivity of reactivation related to the degree of state “negativity”. Instead of averaging across negative/non-negative states, for each trial and each participant, we ascertained the decodability and current value. We then calculated the decodability for each of the 11 values (from -5 to 5, increment by 1), as detailed in Fig. 4B (N=44; error bar = mean ± sem). For negative-value states, we found a significant positive correlation between state value (X-axis) and decodability (Y-axis) (Spearman Rho=0.0860; p= 9.9675e-10; significant signed-rank test above 0 for all 44 sessions’ correlation Rho: p=0.0266), an effect not seen for non-negative states (Spearman Rho=0.0134; p=0.2225; signed-rank test for all sessions’ Rho: p=0.8886). Thus, this finding supports the hypothesis that, during planning, assignment of negative value engenders a weakened state neural reactivation.

We next examined neural dynamics for within-and across-trial negative and positive state neural representations (reactivation decodability). For each trial, and each participant, we down-sampled decodability into 10 trial timepoints (averaged across all 10ms bins within a single timepoint), separately for negative and non-negative states, and then derived their relative decodability (decodability of non-negative states minus decodability of negative states). For each participant, this generated a matrix corresponding to relative decodability of states, within (10 timepoints) and across trials. In a 2-way ANOVA, we found significant mean differences across trials (p<0.0001), but not within, trials (10 time points; p=0.6749), suggesting relative decodability differed significantly across trials. Thus, consistent with participant behaviour (Fig. 1F), we found an across trial increased tendency for greater decoding of non-negative states (Fig. 4C; Spearman rho = 0.7232, p = 2.6876e-08). Plotting these effects separately, it can be seen that this effect is driven both by an across-trial increase in decodability for non-negative states (Spearman rho = 0.7508, p<0.0001) and a decrease for negative states (Spearman rho = -0.6636, p<0.0001) (Fig. S2).

To probe this further, and rule out confounds, we tested if decodability is contributed to by a recency effect, reflecting what subjects had seen in a previous trial. Thus, from the previous trial, we compared the decodability of seen and unseen states, averaged across all trials. We found no significant differences (Mann–Whitney U test p= 0.9037), with decodability of the seen group (0.1665±0.0044, mean±std) slightly lower than that of the unseen group (0.1669±0.0066, mean±std) (Fig. S3A). These results outrule an explanation that across-trial decodability reflect the impact of recency effect.

Additionally, we also asked if decodability is contributed to by an overall familiarity with states by taking the cumulative frequency (accumulated across trials) of state visitation. When asking, at the group level, if this related to state decodability, we found correlation coefficients did not differ from zero (mean±std: -0.0179±0.3345; Fig. S3B; Wilcoxon signed-rank test p=0.6076). Thus, we found no support for an explanation that a relative difference in decodability related to an overall familiarity effect.

To ascertain if value-modulated neural representations (as in Fig. 4B) emerged across trials, we calculated, for each trial, value-based decodability (Spearman correlation rho of values and decodability, averaged rho across 44 sessions) plotted against trial number (Fig. 4D). A significant positive correlation (Spearman rho = 0.5923, p = 1.1127e-05) was evident, consistent with a value-modulation across trial effect.

Recall our behavioural data (Fig. 1F) provided evidence that participants chose higher value states more frequently as a function of learning. On this basis, we asked if the value of states being decoded also increased across trials. For each trial, we summed the most likely decoded state across all 10ms time points, and also across all sessions. We found an across-trial increase in values (Spearman correlation (Rho=0.6640; p= 3.7040e-07), (Fig. 4E). In 27 out of the 44 sessions (Fig. S4), significant positive Spearman correlations (p<0.05) were evident for trial number and sum values of decoded states.

### Correlations of individuals’ neural dynamics and behaviour during value learning

Thus far, we found neural evidence of learning value estimation and state selection during planning. We next asked if these signatures were sensitive to individual behavioural differences. For value estimation, we tested if session performance (averaged final outcome reward value across trials; as shown in Fig. 1C) correlated with a neural value encoding signals (GLM coefficients from the mixed-effect model) (Fig. 5A). The correlation coefficients of these 10 time points were significantly above 0 with signed-rank test (p=0.0488; inset of Fig. 5A), and bootstrap testing controlling for potential autocorrelation of time points (p=0.0144; using the first-order autoregressive integrated moving average (ARIMA) model and bootstrapping 5000 replicates simulating the null hypothesis of intercept=0). We also found a weak positive correlation (Pearson rho =0.1067, p =0.0253; Spearman rho =0.0720, p =0.1319) when we pooled all time points together and controlled for time points using partial correlations. Thus, these results suggest that stronger value-encoding neural signals are linked to better individual performance.

**Figure 5.**
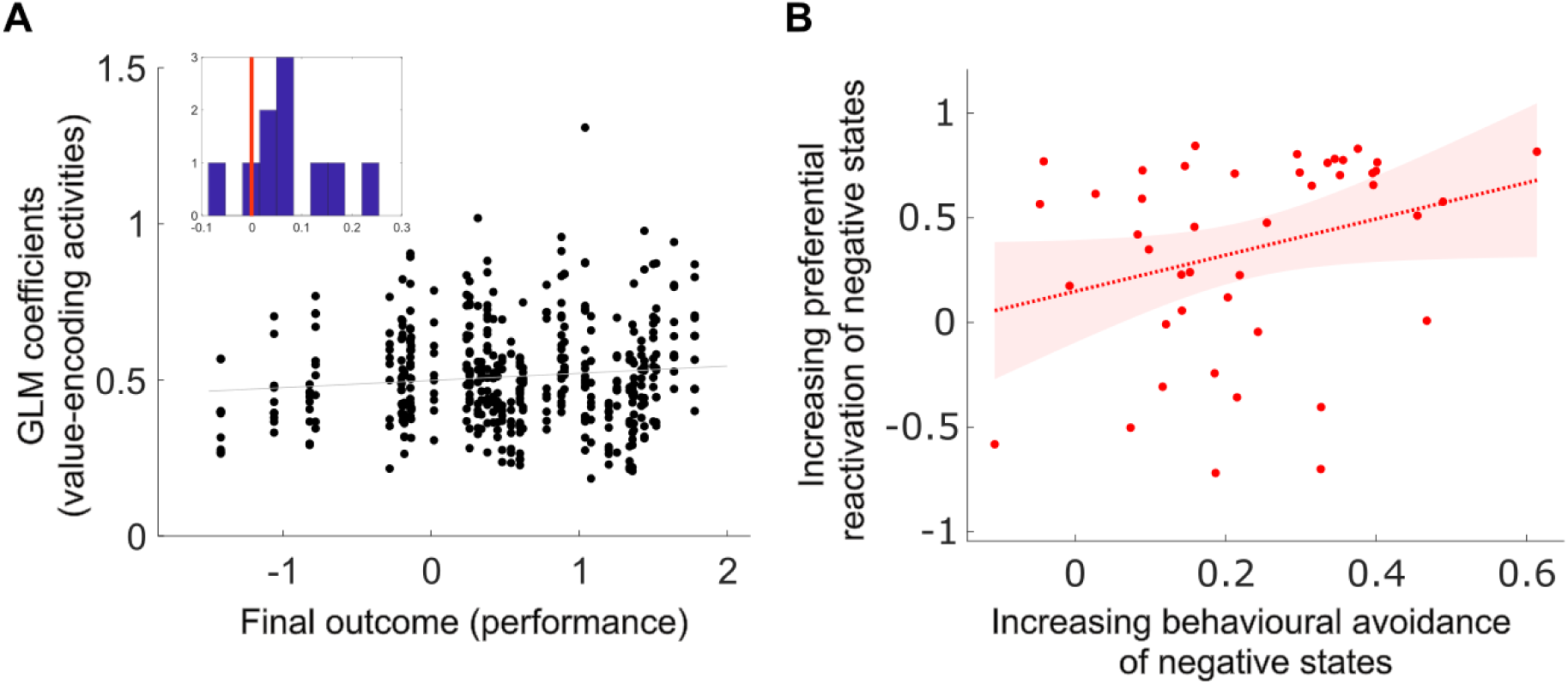
MEG activity and individual differences in value learning. (A) Scatter plot showing the value-encoding GLM coefficients (Y-axis) and task performance of participants (final reward outcome; X-axis), where a weak correlation was evident (Pearson rho =0.1067, p =0.0253; Spearman rho =0.0720, p =0.1319; partial correlation controlled for time points). GLMs’ coefficients represent how much the MEG activity predicts a final outcome value for a participant on each trial. In each of the 44 sessions, a GLM was estimated for each of the 10 within-trial time points, and its coefficients were plotted as solid dots. The inset shows a histogram of all Spearman correlations of the GLM coefficients and the task performance of each of the 10 time points, which are significant positive with signed-rank test against 0 (p=0.0488) and with bootstrap test controlling for potential autocorrelation of time points (p=0.0144; using first-order autoregressive process of ARIMA model, bootstrapped with 5000 replicates simulating the null hypothesis of intercept=0). (B) Scatter plot showing a significant Spearman correlation (Spearman rho =0.3364, p =0.0261, N=44) of individuals’ overall across-trial evolution of behavioural avoidance of negative state options, and the across-trial evolution of selective reactivation of negative states (as shown in Fig. S1 and S4, respectively). The red line represents the least-squares line of all the dots and the shaded region represents the 95% confidence interval.

Finally, we tested whether neural dynamics of state-decodability related to behavioural state selection. We found the degree of across-trial behavioural state disregard (Spearman Rho in Fig. S1; indicating individual sessions’ across-trial discarding of low-value states) correlated with the degree of selective reactivation (Spearman Rho in Fig. S4; indicating the individual sessions’ across-trial discarding of low-value state decodability). Specifically, a significant positive correlation was evident (Fig. 5B; Spearman rho =0.3364, p =0.0261), consistent with the notion of a positive linkage between neural and behavioural avoidance of negative states. These results indicate that individuals’ value encoding neural signals, and selective reactivation of state neural representations (decodability of states) are linked to both individuals’ task performance and a behavioural avoidance of negative states, respectively.

### Theoretical accounts of multi-step planning

To provide a potential theoretical explanation of our findings, we ran simulations using a tree search model previously found to provide a best fit for similar multi-step planning data (Kurth-Nelson et al., 2016) (Fig. 6) that allowed drift in the value for each of the 6 states by - 1, 0, or 1, as well as a random selection of two “neg-flip” states in a trial. This modelling indicated that an RL agent with a tree search policy will exhibit an across-trial increase in final outcome value (Fig. 6A), while also decreasing its choice selection (visit frequency) for low-value or negative states (Fig. 6B & 6C). Thus, our findings in human subjects share core qualitative characteristics to those of an RL agent, where both show an across-trial increase in value encoding signals coupled to a decrease in negative (low-value) state reactivation.

**Figure 6.**
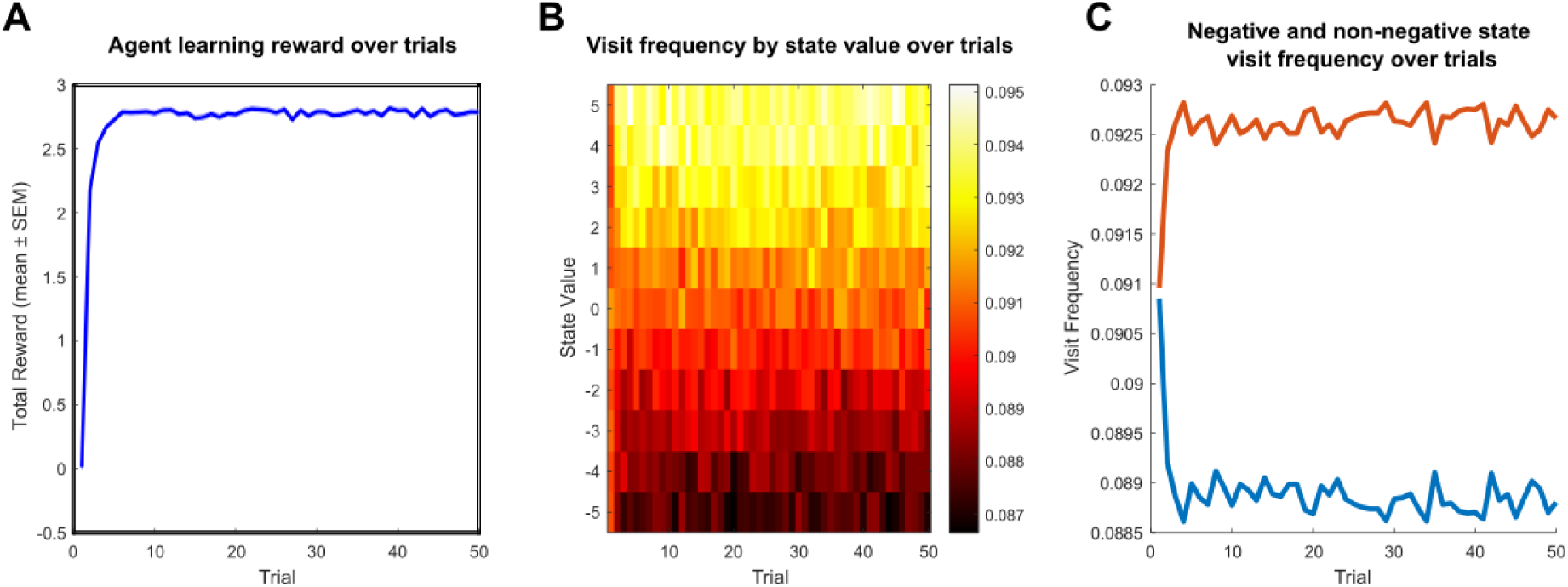
Modelling how RL agents learn multi-step action plans across trials. We simulated the agents’ actions with a decision tree search policy (algorithm), as previously used in multi-step planning research (Kurth-Nelson et al., 2016). We run 100,000 permutated simulations for each of the policies. In every trial in the simulation, the agent needed to perform a 4-step decision task to maximise their overall reward accrual. Similar to the human task, in each trial, the agent started at a random state, and the value of each of the 6 states drifted by -1, 0, or 1. There were also two randomly selected “neg-flip” states in a trial, where the accumulative reward would flip sign (from positive to negative or vice versa) when the agent chose it. (A) Line charts showing the total reward agents obtain across the 50 trials (X-axis) with the tree search policy. Blue lines represent the total reward agents got for that trial, averaged across permutations (thin lines represent the SEM). (B) Heatmaps showing how the frequency of visits changed across trials according to the value of states. In the colour map, brighter colour represents higher visit frequency, and darker colour represents lower visit frequency. (C) Line charts showing the averaged visit frequency across trials, for the negative (blue lines) and non-negative states (red lines).

## Discussion

Learning to plan involves adopting a strategy (policy) akin to that of reinforcement learning (RL) agents, learning values across trials, selecting policies according to need, and availing of flexible model-based and inflexible model-free control (Gershman, 2018; Venditto et al., 2024). In its simplest terms, agents learn to estimate value and choose actions accordingly (Sutton & Barto, 1998). Here, we found that humans share dynamics with RL agents in how they learn to choose states with higher values across trials.

In a previous related work, involving a similar task structure (Kurth-Nelson et al., 2016), we found that most participants showed model-based tree-search-like behaviour. Consequently, the modelling we adopted in this study focused solely on such tree search. Note that our study is not designed to distinguish which policies are represented in the multi-step planning, and we cannot exclude the possibility that participants used other policies. Future research, using bespoke designs, will need to address whether agents develop and integrate different policies during planning. In our research, we focused on decoding sequential state reactivation (replay) (Liu et al., 2019, 2021; McFadyen et al., 2023; Schwartenbeck et al., 2023) and here we extend on this to ask how the dynamics of neural representations relate to the dynamics of planning. Here, we build on a suggestion (Hunt et al., 2021) that investigating neural state representations is critical for understanding how we plan in dynamic environments.

A striking finding in our study was that neural state reactivations, as determined by their decodability, depend on their values, with these also reflecting an evolving across-trial learning process. Rodent studies provide evidence for differential value-modulated reactivation in hippocampal place cells, with stronger reactivation and replay for high reward locations (Singer & Frank, 2009; Ambrose et al., 2016; Michon et al., 2021; Jin et al., 2024). Consistent with this, using out-of-sample decoders, we show that human state reactivations also relate to value, including an evolving across-trial relative decrease in decodability for states with negative values. We note that, in rodents, there is enhanced place cell remapping when animals experience negative (aversive) events (Moita et al., 2004; Ormond et al., 2023; Blair et al., 2023). A decreased decodability for negative states in our study is likely to reflect this greater susceptibility to remapping, rendering corresponding state reactivations less decodable.

A key finding was an emergence of stronger and earlier value encoding signals following learning, echoing a shift in dopamine prediction error signals from unconditioned (US; unexpected reward) to conditioned stimuli (CS; cue onset) following learning, as in temporal difference (TD) models (Amo et al., 2022). At the beginning of our experiment, participants cannot accurately estimate how much reward (or state value) they will get, but learnt this across trials, as evident in the emergence of an earlier planning value-encoding signal as predicted within an RL-TD framework.

In computational neuroscience, recent work has extended traditional drift-diffusion models (DDM) (Ratcliff 1978; Ratcliff et al., 2016; Prat-Ortega et al., 2021) into RL-based evidence accumulation models (EAMs) to explain across-trial learning in decision making (Pedersen et al., 2017; Fontanesi et al., 2019; Miletić et al., 2021). While these models detail how neural dynamics contribute to single choices based on a decision threshold, but are less adept at explaining multi-step planning. A more recent computer science approach utilises recurrent neural network (RNN) models to explain multi-step behaviour (Zhang et al., 2020; Ji-An et al., 2025) as well as within-trial neural mechanisms (Mante et al., 2013; Yang et al., 2019; Rajalingham et al., 2025). In these RNNs, hidden units and output units accumulate evidence from an input until a confidence threshold (or other decision criterion) is met. RL-based recurrent neural network (RNN) models have also been adopted to investigate value-guided behaviour of humans and animals (Eckstein et al., 2023; Miller et al., 2023; Ji-An et al., 2025). Future studies might usefully test if this could be a plausible model in explaining our finding of earlier and stronger value encoding signals (accumulative evidence) with across-trial learning.

Although precise anatomical conclusions cannot be drawn with MEG, using source reconstruction, we found boosted vmPFC activity with the evolution of value learning (Fig. 4F). Interestingly, a previous human MEG study (Hunt et al., 2012) found the very opposite, reporting a decrease in vmPFC activity across trials. A possible explanation for vmPFC being more actively engaged in our study relates to the need to compute multi-step decisions, wherein subjects are tasked to remember state values and engage in active computation as to the likely outcome of a plan. In the aforementioned study (Hunt et al., 2012), such a complex calculation and revaluation is not required. We also note that previous research implicates other brain regions, such as the orbitofrontal cortex (OFC), ventral striatum, and midbrain, in value estimation and reward learning (Schultz et al., 1997; Dolan & Dayan, 2013; Massi et al., 2018; Ballesta et al., 2020; Knudsen & Wallis, 2022; Gore et al., 2023; Ferrari-Toniolo & Schultz, 2023; Yun et al., 2023). We did not find across-trial signal changes in these regions using a standard threshold (Z>3.1), but this most likely reflects the low sensitivity of MEG for detecting activity in deeper regions (Hillebrand & Barnes, 2002).

GLMs are widely used in the analysis of human fMRI and animal neurophysiological studies (Friston et al., 1994; Pillow et al., 2008; Muhle-Karbe et al., 2023), while in MEG and EEG studies, evoked (event-related) potentials (ERP) and time-frequency analysis are more frequently used (David et al., 2006; Puce & Hämäläinen, 2017). Our use of GLMs enabled us to account for diverse reward amounts and planning time in value estimation, facilitating a probing of underlying learning-related neural dynamics. Utilising tools that align dynamic temporal responses, such as dynamic time wrapping techniques (Berndt & Clifford, 1994; Williams et al., 2020), might be usefully availed of in the future to investigate ERP or frequency dynamics related to value encoding. As previous research probed how ERP and frequency power (mainly theta and beta) dynamics evolve during learning (Cavanagh et al., 2010; Van de Vijver et al., 2011, 2018), including revealing greater theta power linked to stronger state reactivations (Wise et al., 2021), this suggest a valuable direction for future studies might be to examine the precise connections between frequency power and the types of evolving state and value-encoding dynamics we detail in the current study.

In conclusion, by tracking the dynamics of neural signals, within and across trials, we show an emergence of stronger and earlier value encoding signals, linked to boosted vmPFC activity. Crucially, we show that neural reactivations (decodability) are value-dependent and linked to state value learning across trials. These results connect behavioural, neural, and theoretical accounts of learning multi-step decisions, establishing a foundation to investigate how healthy individuals and those with disorders putatively linked to distorted RL patterns (Pike & Robinson, 2022) learn to solve multi-step problems.

## Methods and Materials

### Participants

We recruited 44 sessions from 22 adults (17 females and 5 males) aged 19-28 years old (22.9545±0. 0.5439, mean ± sem) from the subject pool of the University College London (UCL) Institute of Cognitive Neuroscience. All participants reported an absence of a history of psychiatric or neurological disorders. All participants provided written consent, including consent to publish their data. The experiment was approved (ethics number 1825/005) by the Research Ethics Committee at UCL in the UK.

### Task

We availed of a previously unpublished data set collected originally as a replication study of Kurth-Nelson et al., 2016. Our focus here was solely on elucidating neural dynamics related to multistep value planning. This task was implemented in Matlab (MathWorks), where in each trial participants were required to perform a 4-step decision task to collect as much reward as possible (Fig. 1). Each state is associated with a unique visual stimulus (6 objects pseudorandomly drawn from a set of ten subjects: bird, bread, cat, chair, garlic, hammer, hand, horn, tree, water) and a state value (between -5 and +5 pence) that drifted on each trial pseudorandomly by -1, 0 +1. The trial started with an initial state that was psedorandomly selected. From each of the states, the participant could move “up” or “down” by pressing a button, which would deterministically lead to another state. After reaching a new state, the state value would be added to the total accumulated reward they got. To discourage responses without planning, two “neg-flip” states were psedorandomly selected for each trial. If they reach the “neg-flip” state, the sign of the total cumulative reward earnings until (including) that state would be flipped (e.g. 9 became -9; -6 became 6). Before the scanning, the participant learned the transition structure of the task but not the initial state value.

In each trial, we showed the participants the starting state and two “neg-flip” states in text. Then they would have up to 60s (self-paced) to plan, and they could stop planning by pressing a button. After the planning period, they would be shown a blank screen and would have to enter the 4 moves (up to 3s to enter the first and 1s for each of the other three moves). Participants were then asked to repeat the moves (up to 10s were allowed to execute a move) they entered when the visual stimuli were shown one by one, corresponding to the transition being entered. Each execution of move was followed by a cross-fade transition (350ms), a 500ms pause, 1000ms for displaying the reward amount of the current state, 1000ms for displaying the total running cumulative earnings until then, and finally, if any, a negative change to the total trial earnings. If this was the final move, the final reward they got for the trial would be shown for 3000ms; if not, then they could enter the next move. Only one state was shown to the participant at once, and no bird-eye overall view was ever shown. About 50 trials (dependent on the scanning time available) were collected from each participant.

We calculated the performance (reward outcome) based on the “believed reward”, where the value of stimuli is the value that would pertain if the states had not changed since the last time they were encountered, which is identical to having a 1.0 learning rate with planning. This approach captures the value participants believed they would get, instead of a value with added in random noise by virtue of a drifting state value. We used the final total value (sum of these believed rewards and flipped if there was any “neg-flip” state involved) to determine the performance of participants.

After the 6-state navigation task, a localiser task was presented to the participants while they were still in the scanner. Similar to the previous study (Kurth-Nelson et al., 2016), this task was used to train classification (Lasso-regularised logistic regression) models. During the task, the name (text) of a visual stimulus appeared on the screen for 1500-3000ms (varying). Then the visual object itself was shown. In 20% of the trials, the visual stimuli were turned upside-down. The participants were asked to press a button when the stimulus shown in that trial was correct side up, and press another button when it was not. A green or red fixation cross was then shown, respectively, to indicate if they answered it correctly after the response. After that, there was a 700-1700ms inter-trial interval (varying). In this localiser task, a total of 125 pseudorandomised trials (about 16 correct side up trials for each stimulus, which were used for the model training) were collected from each participant, with the same 6 objects used in the main task.

### MEG data acquisition

MEG data was recorded with a CTF Omega 275-channel axial gradiometer system, with a 1200 samples/second sampling rate. All data were collected in the Wellcome Centre for Human Neuroimaging of the UCL Queen Square Institute of Neurology. Participants’ responses were recorded with a button box, and they were asked to put their fingers in the position they felt most comfortable with.

All data was downsampled to 100Hz to improve signal-to-noise ratio, and further filtered with a 0.5Hz high-pass filter, then removed artefacts with an automatic Independent Component Analysis (ICA) filtering signals that matched the EOG channels. All analyses performed here were based on the filtered data.

### GLM analyses

To account for the diverse planning time across trials and subjects, we normalised the planning time to 10 time points and further built GLM models based on these normalised data. To study the dynamic change across trials (following value learning), the trials were divided into 4 quarters. We first test if the value encoding signals emerged after learning: for each quarter, we built a GLM (using sensor-specific activity to predict final outcome value) for each sensor within each participant, illustrated that in the heatmap (Fig. 2A), and tested with a 2-way ANOVA test (tested for within and across trials dynamics, with subject ID as a control). Then we took the absolute coefficients and averaged across all sensors (Fig. 2B). We ran a Spearman rank correlation to test if the signal increased across the quarters.

To further investigate the interaction of within and across trial dynamics, we adapted a mixed-effect GLM model. We put in the final outcome values, trial number and their interactions as regressors to predict the MEG activity (magnitude), therefore building an encoding model to index the contribution of these variables to the activity. After that, we built group-level GLMs using the first-level coefficients (used as dependent variables) and the subject identity as dummy variables to regress out the subject-specific signals. In this mixed effect model, we found that the GLM coefficients of the interaction of trial numbers and values decreased across the 10 planning time points using Spearman rank correlation (Fig. 2C).

### Decoding analyses

One feature of value-based learning is the discarding of negative states (Huys et al., 2012; Faulkner et al., 2021). We therefore tested whether the neural representations of negative states were discarded and if that was modulated by how negative these states were. We trained decoders with an out-of-sample localizer at the end of the task. For each participant, we trained a Lasso logistic regression model at 200ms after visual stimuli onset as in previous studies (Kurth-Nelson et al., 2016; Liu et al, 2019), to predict the neural representations of the six states. To verify the decoders, we tested the accuracy and predicted probability of each state and compared that with the baseline (1/6 or 16.67%) (Fig. 3), where the accuracy was calculated based on if the predicted label (the state with the highest predicted probability at a particular time point) matched the actual label. These trained decoders were then applied to the planning phase MEG data, in which no visual stimuli were shown. We predicted the neural representations (one of the six states) based on the highest predicted probability. Then, we calculated the percentage of each state being predicted (decodability) within a trial and averaged these percentages across the states with negative values and non-negative values separately. Then, we separately averaged these two percentages across trials for each participant (Fig. 4A). Next, we tested if the avoidance of the negative state representations was modulated by the values (the “negativity’) of these states. For each trial in each participant, we calculated the decodability and its current value. We then calculated, for each of the 44 sessions, the decodability (neural representations) of each of the 11 values (ranging from -5 to 5, increment by 1; Fig. 4B). We correlated the neural representations (Y-axis) with the value of states (X-axis) using Spearman rank correlation to test if there was any modulation effect.

We also further tested if these discarding and modulations were learned across trials. We calculated, for each trial, the representations differences of negative and non-negative states (averaged across participants) and Spearman rank correlated that with the trial number (Fig. 4C). Moreover, to test if the value-modulation effect was learned across trials, we Spearman rank correlated the trial number with the trial-by-trial Spearman Rhos (slope of Fig. 4B; i.e. correlations of state values and the neural representations). With these, we verified our hypothesis that these selective reactivation features are learned across trials.

As described in previous studies (Huys et al., 2012; Faulkner et al., 2021), subjects may discard states with low/negative values during planning. Therefore, we tested here whether participants discard low-value states in their action planning, in terms of both behaviour and neural representations (reactivations).

For the behavioural analyses, for each trial, we summed the values (believed values, assuming no changes of state values since that participant saw it last time) of the 4 states that one participant would go to (according to what one entered), and correlated that with the current trial numbers with Spearman rank correlation. We repeatedly performed the analyses for each participant and also across all participants (Fig. 1D; Fig. S1). We performed a similar analysis for the neural presentation analysis: for each trial in a participant, we calculated the value based on the multiplication of the state value and decodability of that state during the whole planning phase (decoded using an out-of-sample localizer). Similar to the behavioural analysis, we performed Spearman correlation using the value and the trial numbers within and across participants (Fig. 4E; Fig. S4).

### Source reconstruction

We investigated the underlying neural source changes following value learning using beamforming source localisation. We epoched the MEG data using a -100ms to 2000 ms window and transformed the data into a 2mm MNI space with a linearly constrained minimum variance beamformer in OSL, as in previous studies (Van Veen et al., 1997; Mattout et al., 2006; Liu et al, 2019; Wimmer et al., 2020). We generated a single shell with superposition corresponding to the MEG sensors’ plane tangential. The sensor covariances were estimated using a general frequency range (1-45 Hz), as used in the previous study (Wimmer et al., 2020). We performed 5000 non-parametric permutations in OSL, calculated t-maps and smoothed with a 2 mm FWHM Gaussian kernel. We then performed a paired t-test comparing the first and the last block of each session, as a group-level analysis, when applying an independent vmPFC mask (Delgado et al., 2016; Bhanji et al., 2019) to the data and calculating the statistics (SVC; P<0.05, corrected for FWE).

### Individual analysis

Individual analysis was performed to test if the neural signatures we found can be related to individual behaviour. For value estimation, we tested if the value encoding signals (GLM coefficients) of individual participants were linked to how much one could earn across the trials. For each of the 10 planning time points, we correlated the GLM coefficients of the value regressor from the mixed effect model with the amount of final total value (assuming state values have not been changed since the last time they saw it). We correlated these two parameters using Spearman rank correlations (Fig. 5A). Then, we tested across time points to see if these Spearman rank correlation Rhos were significant above (or below) zero using a Mann–Whitney U (ranksum) test (Fig. 5A inset).

For action planning, we tested if the behavioural avoidance of low-value states (as seen in Fig. 1D & Fig. S1; indicating the learning of discarding the low-value state actions) would be correlated with the neural representational disregard of low-value states (as seen in Fig. 4E & Fig. S4; indicating the learning of discarding the low-value state neural representations).

These were plotted in the scatter plot and correlated with Spearman rank correlation (Fig. 5B).

### Simulations of multi-step planning

We implemented reinforcement learning agents that used the full-depth model-based tree search policy, to solve the multi-step planning problem we asked the human participants to solve. This model-based tree search agents (Sutton & Barto, 1998; Gershman, 2018) employed an exhaustive forward planning strategy with depth d=4, identical to the number of actions. We employed this exhaustive strategy because this model fit well with human behaviour (Kurth-Nelson et al., 2016).

For each of the agents, we ran 100,000 independent simulations, each containing 50 trials of the 4-step decision task. Similar to the human participants, the value of the states was unknown at the beginning of a simulation, and the agents needed to learn the value of states across trials. In each trial, they started at a random state and needed to make 4 moves to maximise the reward they got. At each trial, the state value of the 6 states drifted by -1, 0, or 1; and there were also two randomly selected “neg-flip” states, where the cumulative reward the agents got until that point would flip sign (from positive to negative or vice versa) when the agent chose it.

In this model, the agents made choices to maximise the value one gets from all possible paths based on the following value function equation, with the value of states learned across trials:

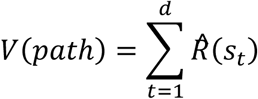

Where t is the depth of the step trajectory, and 𝑠_𝑡_ represents the state at depth t. The estimated reward for each state𝑅̂(𝑠_𝑡_) is updated at the end of each trial through the following equation:

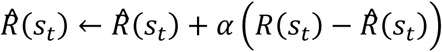

Where 𝛼 represents the learning rate, and 𝛼 =0.9 was used in the simulation.

## Acknowledgments

This work is supported by the Max Planck UCL Centre, a joint initiative supported by the Max Planck Society and UCL. We acknowledge funding from the Max Planck Society to R.J.D. (549771-D.CON 177814) and a Wellcome Trust Investigator Award to R.J.D. (098362/Z/12/Z);

We would like to thank Yunzhe Liu for data collection; Zeb Kurth-Nelson for helpful suggestions and comments; Rani Moran, Lennart Luettgau, Matthew Nour, Veith Weilnhammer, Pierre Vassiliadis, Maria Eckstein, G. Elliott Wimmer, Mehrdad Salmasi, and Tricia Seow for comments.

## Supplementary

**Figure S1.**
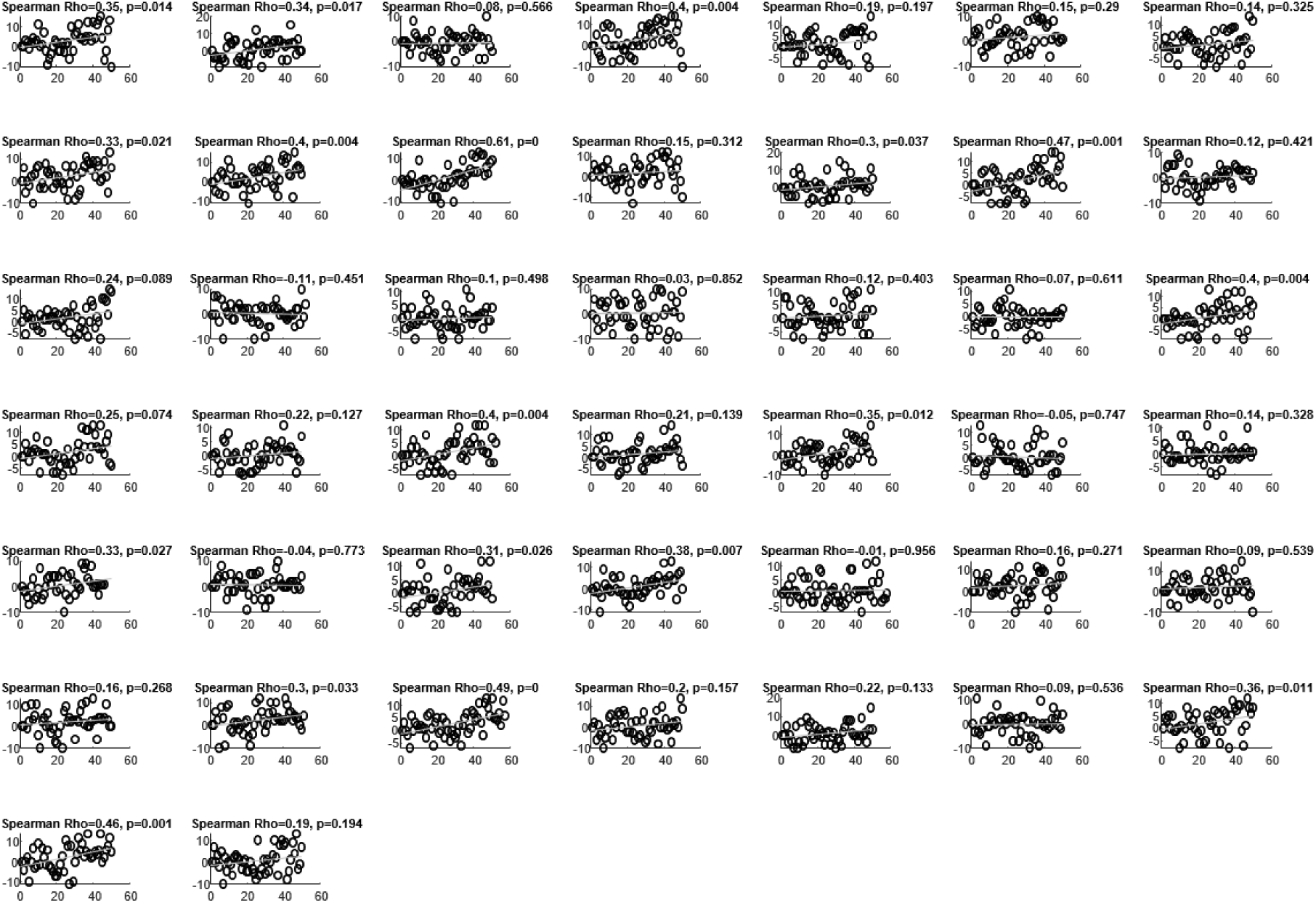
Correlation of individual participants’ trial number (X-axis) and the sum of the chosen state values (value they would get if the states have not been changed since the last time they saw it) (Y-axis).

**Figure S2.**
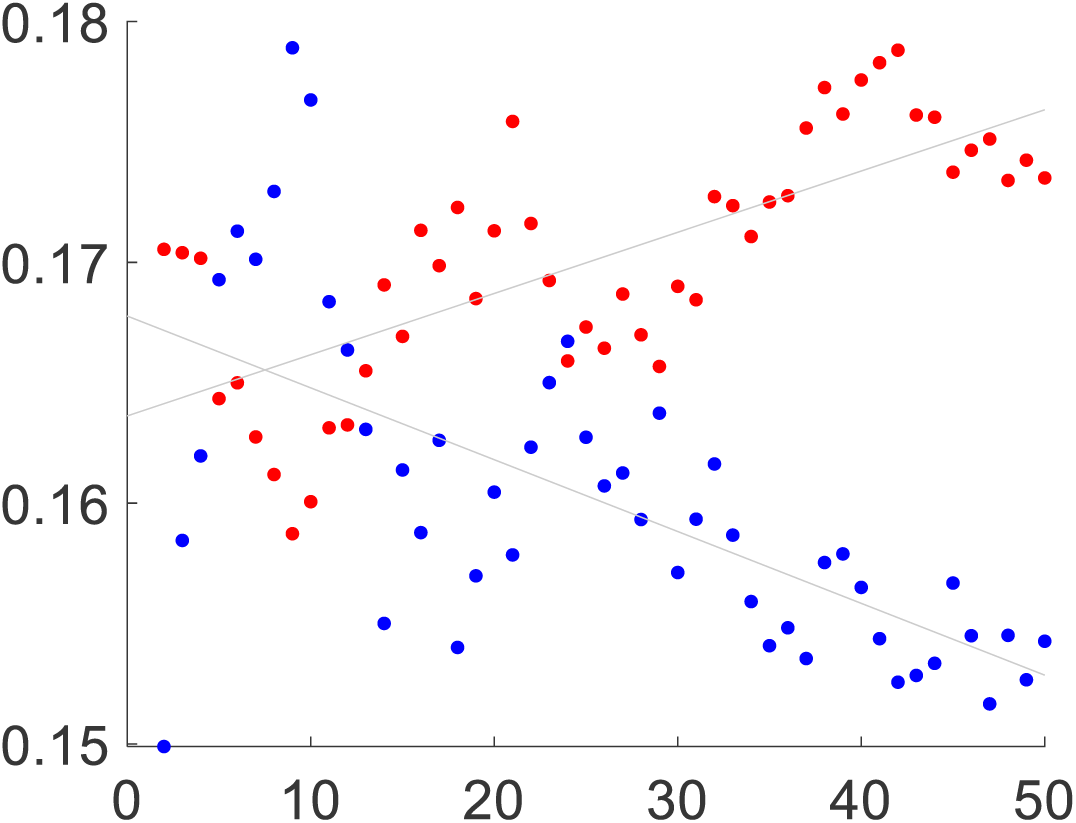
Scatter plot showing the correlations of decodability (Y-axis) and trial number (X-axis), for negative (blue) and non-negative states (red).

**Figure S3.**
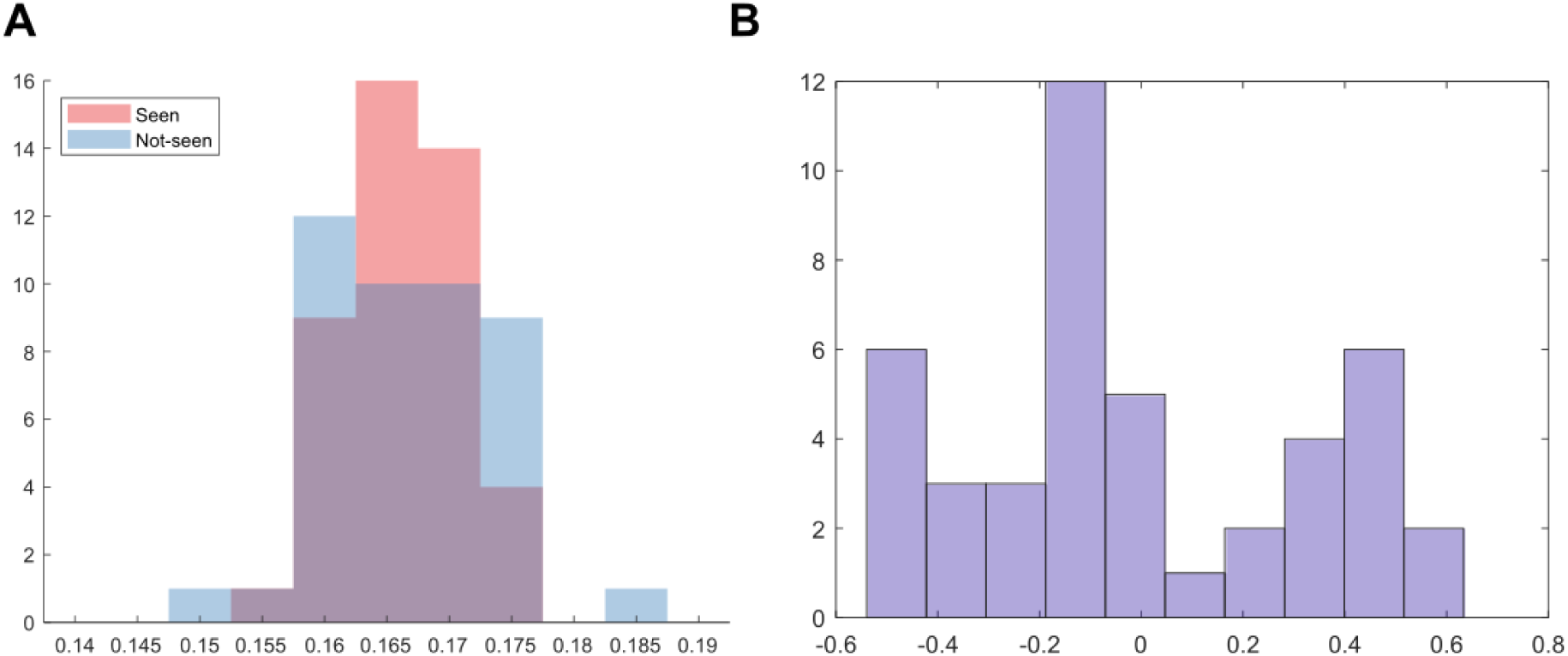
Histogram of recency effect and familiarity effect tests. (A) Distribution of decodability in seen and not-seen (in the last trial) groups, as a test for recency effect. (B) Distribution of Spearman Rho for correlations of state visit frequency and its decodability, as a test for familiarity effect.

**Figure S4.**
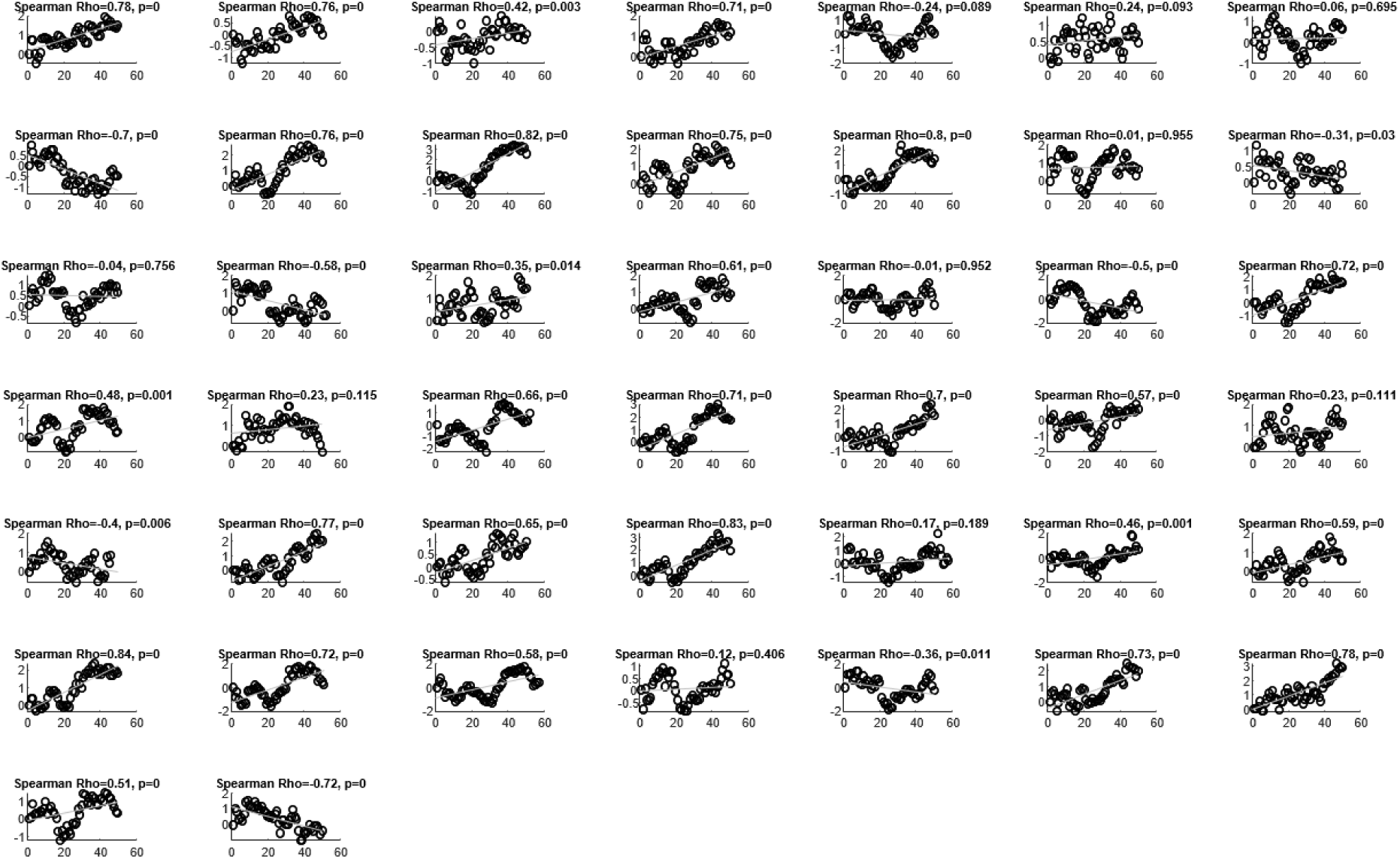
Correlation of individual participants’ trial number (X-axis) and averaged values of the neural representations of states (Y-axis).

